# NeuroCaptain v2 – Interactive Three-Dimensional fNIRS Optode and Probe Montage Design Platform Based on Blender

**DOI:** 10.64898/2026.06.02.729322

**Authors:** Ashlyn McCann, Qianqian Fang

## Abstract

**Significance:** Accurate and reproducible optode placement is crucial for obtaining high-quality fNIRS data in both individual and group-level neuroimaging studies. Conventional optode/probe montage design tools usually transform a probe layout defined in 2D Cartesian space onto a 3D head surface using a mass-spring model. Such mechanical transformation, combined with the indirect mapping between the 2D probe definition and the 3D target space, can introduce placement variations across different head surfaces and subjects.

**Aim:** We introduce NeuroCaptain v2, an open-source Blender-based add-on designed to enable interactive, anatomically guided optode design, registration, and cortical sensitivity visualization for fNIRS head-cap and probe creation.

**Approach:** NeuroCaptain v2 enables researchers to add, move, and edit fNIRS sources and detectors directly over a 3D head surface mesh, defining anchored optode positions, as well as setting the stiffness of the springs between adjacent optodes. It then utilizes Blender’s built-in physical simulation engine to relax the initial probe layout to satisfy the mechanical constraints. With the built-in mesh-based Monte Carlo (MMC) and diffusion-solver Redbird, NeuroCaptain v2 computes and renders 3D sensitivity maps to guide iterative optode adjustment. The resulting 3D optode layout is stored in the form of barycentric coordinates defined in a 10-20 landmark mesh, enabling consistent probe transfer across different head models.

**Results:** We demonstrate interactive 3D montage design, cross-head-atlas probe registration, and cortical sensitivity visualization across multiple head geometries. Registering a probe across seven neurodevelopmental head atlases, the proposed anatomical-coordinate approach yields a mean per-optode standard deviation of 2.29 mm, a roughly 74% reduction in cross-subject placement variability compared to 8.68 mm using a conventional 2D-to-3D registration.

**Conclusions:** NeuroCaptain v2 provides a reproducible, fully open-source workflow for fNIRS probe montage design that facilitates anatomically guided probe development and cross-subject registration directly in a three-dimensional anatomical environment.

## 1 Introduction

Functional near-infrared spectroscopy (fNIRS) has emerged as a versatile neuroimaging modality for both research and clinical applications.^1^ An fNIRS system typically consists of an array of low-power near-infrared (NIR) light sources and detectors, known as the optodes, secured over the subject’s head using a head-cap. The wearability of lightweight fNIRS head-gears and safety for long-term measurement make fNIRS an attractive option in applications outside of the restrictive environments traditionally required for functional magnetic resonance imaging (fMRI) based measurements.^2^ Recent advances in modular circuitry design, compact light-emitting-diode (LED) light sources, and high performance light detectors have enabled researchers to build high-density, high-performance wearable fNIRS probes,^3, 4^ further unlocking the potential of monitoring brain activities in day-to-day tasks.^5, 6^ Obtaining high-quality and robust fNIRS measurements in such complex and dynamic tasks is crucial for advancing our understanding of human brain function and for monitoring the progression of neurological diseases.^7^ Designing flexible fNIRS head-gear for increasingly challenging experimental tasks, and creating optode montages capable of capturing the relevant brain activities, are key to further expanding fNIRS research.

There are several well-recognized challenges inherent to fNIRS instrumentation and experimental designs. These challenges are further amplified by the shift toward naturalistic paradigms.^8–10^ One of the known challenges in fNIRS studies is the consistent optode placement across study sessions and subjects,^11^ especially for group analyses.^12, 13^ This issue is particularly pronounced in the emerging hyperscanning studies, where evaluating functional synchrony between multiple participants^14, 15^ relies heavily on consistent optode placement to ensure the desired functional cortical regions are accurately monitored simultaneously. Although inconsistent probe placement between hyperscanning subjects can be partially compensated using post-processing, such compensation often requires additional acquisition of the actual optode locations and, in many cases, subject-specific anatomical scans.^16^ These steps not only require extra equipment and measurement procedures, including digitization and photogrammetry^17, 18^, but the effectiveness of the correction can also depend heavily on the optode layout and the severity of the misalignment.

Designing the fNIRS probe to provide sufficient coverage to the target region-of-interest (ROI) is another known challenge in fNIRS.^19^ This is further exacerbated because complex and dynamic experimental paradigms often elicit broad neural activation across multiple ROIs due to concurrent and diverse stimuli.^20, 21^ Although full-head fNIRS probes are increasingly available, the added system cost and more complex data analysis still limit their widespread use. Accounting for probe coverage and multi-subject anatomical variations during the head-gear and probe montage design stages, in conjunction with additional head-gear placement monitoring methods^22^, could substantially enhance the robustness of the measurement data.

One approach to improving probe coverage and subject-head-shape adaptivity is through custom headgear. NinjaCap,^23^ for example, provides a platform for creating head-size and optode customizable three-dimensional (3D)-printed headgear by translating anatomical coordinates into flat-print templates followed by manual assembly. Our lab previously reported NeuroCaptain,^24^ an open-source Blender add-on that generates anatomically derived 3D-printable head caps with integrated head landmark markers. By building on Blender’s intuitive interface, NeuroCaptain streamlines the cap generation process into an accessible and user-friendly workflow. Despite these advances, both tools are limited to head-cap design and fabrication and do not specifically address probe montage design and optimization. Probe layout, channel configuration, and source-detector registration to the subject head surface require additional software tools, resulting in a fragmented workflow that complicates the integration of probe design with headgear fabrication.

Optode montage design is a foundational step in the fNIRS experimental workflow, and several open-source tools have been developed to facilitate this process. Notably, AtlasViewer^25^ is a widely used MATLAB-based toolbox for designing optode layout in two dimensions and registering them to a three-dimensional anatomical head model using a mass-spring-based relaxation model, providing metrics for evaluating probe performance including inter-subject variability and cortical sensitivity via forward modeling. The fNIRS Optodes Location Decider (fOLD) toolbox^26^ optimizes channel configurations to maximize sensitivity to user-specified cortical ROIs using photon propagation simulation, with the devfOLD^27^ extension broadening this framework to use age-specific atlases^28^ to address developmental fNIRS research. The modular optode configuration analyzer (MOCA)^29^ uses semi-automatic algorithms to tile and interconnect multiple fNIRS modules, such as our lab’s modular optical brain imaging device,^30^ over a user-specified ROI while reporting quantitative metrics including spatial channel distribution and average brain sensitivity. Together, these tools represent significant advances in fNIRS montage design, providing researchers with quantitative frameworks for optimizing channel configurations, evaluating cortical sensitivity, and supporting anatomically informed probe placement across diverse experimental contexts.

To the best of our knowledge, none of the existing optode design software is capable of directly designing, evaluating and optimizing a probe montage in the 3D subject head space. For simplicity, most existing tools iteratively project between a two-dimensional (2D) space where the montage is defined and the 3D atlas/subject head surface. The discrepancy between the optode montage definition space and the physical space where the probe performance metrics are evaluated can complicate the optimization process^29^, underscoring the importance of 3D anatomical information for increasing registration accuracy.^31, 32^ Additionally, most existing montage design tools focus on designing and optimizing a probe layout over a single canonical atlas model. The lack of diverse and subject-specific surface support could lead to suboptimal probe montage designs.

NeuroCaptain v2 addresses several of these unmet needs by extending the previously published NeuroCaptain platform^24^ to support a direct 3D optode montage design workflow. As a Blender add-on, it leverages Blender’s rich 3D modeling and interactive editing capabilities to enable intuitive design and model-guided optimization directly on the 3D head surface, circumventing the limitations of indirect 2D-to-3D dual-space iterations. By incorporating our model-based forward simulators, the mesh-based Monte Carlo (MMC) solver and the diffusion-equation solver Redbird, NeuroCaptain v2 efficiently computes optode sensitivity maps to offer users direct guidance for interactively adjusting optode positions and spacing. Instead of representing the optode positions in a 2D Cartesian space, NeuroCaptain v2 utilizes anatomical coordinates to store and transfer a designed montage from one head model to another.

In the following Methods section, we first describe the key technical advances implemented in the NeuroCaptain v2 workflow, including the interactive 3D optode montage design process, sensitivity-informed optode optimization, anatomical-coordinate-based optode definitions, registration between subjects, and integration with the 3D-printable head-cap design workflow. The effectiveness of these technical approaches is evaluated in the Results section using sample probe layouts; the repeatability of registering a single probe and of registering across different head surface models is evaluated and compared against an established montage design tool. Finally, we discuss implications of NeuroCaptain v2 for standardized fNIRS probe design as well as its current limitations and future work.

## 2 Methods

### 2.1 NeuroCaptain v2 Workflow Overview

NeuroCaptain v2 offers fNIRS researchers an intuitive and interactive platform for directly designing, editing, and optimizing complex optode/electrode montages over any 3D head surface model. By replacing the previously used 2D Cartesian representation of the optode/probe definition with anatomical coordinates in the form of barycentric coordinates within a tessellated 10-20 landmark mesh, NeuroCaptain v2 directly links probe design requirements (such as anchor positions and spring properties) with the 3D head space where the probe sensitivity is computed and coverage is optimized.

The NeuroCaptain optode montage design workflow is illustrated in the left side of Fig. 1. It begins with a head surface mesh, from which the head 10-20 landmarks are automatically computed.^24^ In contrast to other existing probe design tools where only a canonical atlas is used, NeuroCaptain can dynamically download pregenerated head models from several dozen Neurodevelopmental MRI atlases hosted on our NeuroJSON.io data portal.^33^ Using the toolbar provided in the NeuroCaptain add-on, users can conveniently place sources and detectors over the head surface to form an initial probe montage. NeuroCaptain automatically tessellates the sources and detectors to build the edges connecting adjacent optodes. Subsequently, users can interactively select any optode and define it as an anchor – an optode held at a fixed position during the following spring-relaxation step; users can also select each edge between optodes and set its individual stiffness, including rigid springs (high stiffness). By invoking Blender’s built-in physical simulation engine, a spring relaxation simulation is performed. Once the steady state is reached, the 10-20 barycentric coordinates of the final optode positions are updated and exported, if requested, in a plain-text JavaScript Object Notation (JSON) file.

**Fig 1:**
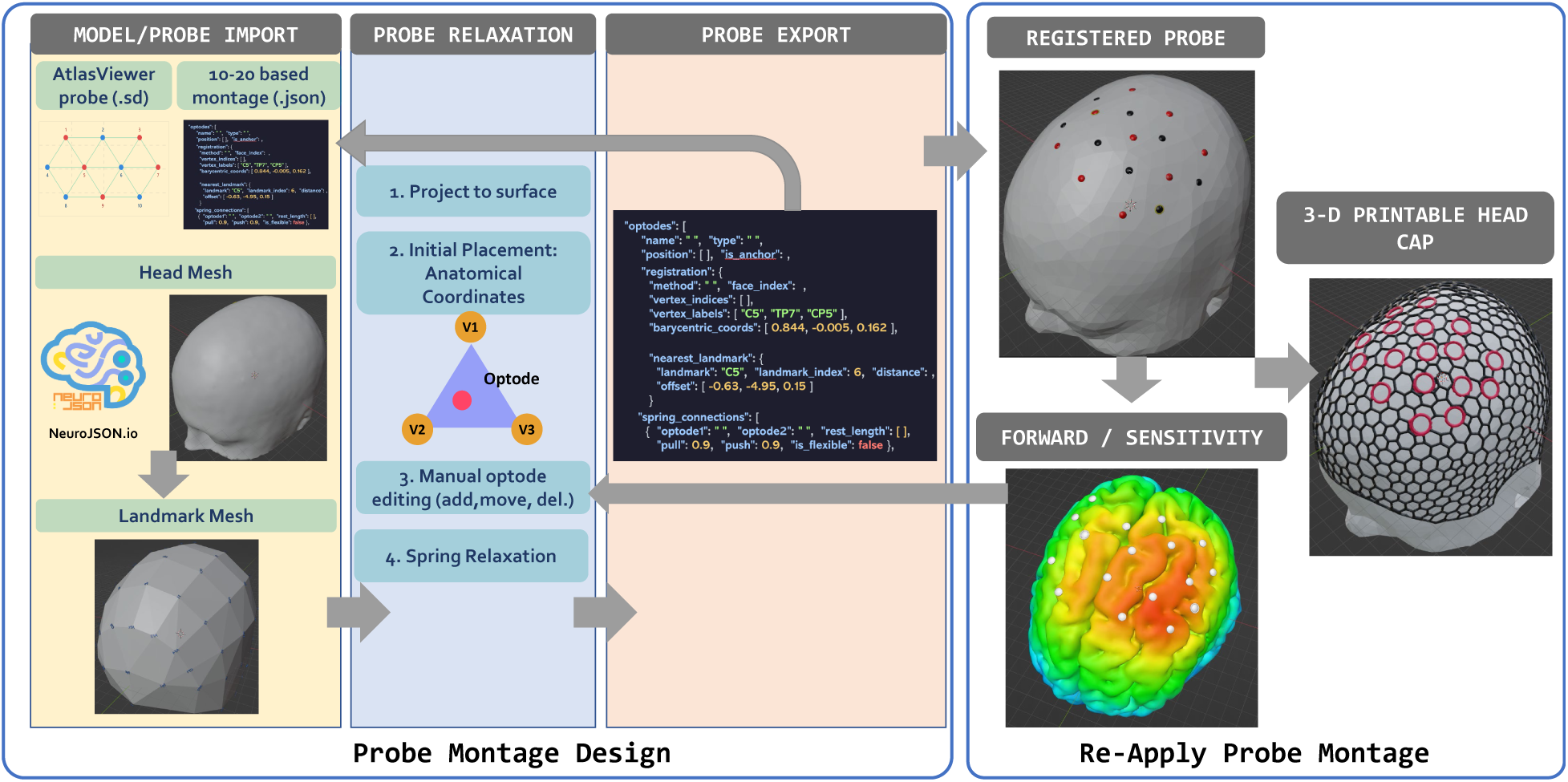
Overview of the NeuroCaptain v2 workflow. The left-panel shows the probe/montage design workflow. The user inputs (optode JSON configuration, head mesh, and landmark mesh) are processed through three registration stages: surface projection, barycentric initial placement, and spring relaxation. The right-panel depict steps for re-applying a saved probe to a new head surface – including spring relaxation to register the probe, compute cortical sensitivity map to guide the manual adjustment of the optodes, and finally creating optode-embedding 3D printable head caps.

The process for loading and registering a saved montage design over a new head surface model is depicted on the right half of Fig. 1. Because the optodes are represented in the 10-20 based anatomical coordinates, all optodes are initialized according to the 10-20 landmarks of the new head surface. To simultaneously satisfy the mechanical constraints of the mass-spring model, a second relaxation is performed to warp the probe to the new head surface. Next, NeuroCaptain allows users to choose either MMC or Redbird to perform forward simulations across the designed montage and render the resulting 3D sensitivity map on the cortical surface. Based on the sensitivity distribution, users can move the optode positions and rerun the relaxation and sensitivity computation. Finally, the optode positions can be integrated as landmarks into the wireframe 3D-printable head-cap following the workflow reported in our initial NeuroCaptain publication.^24^

### 2.2 Head Mesh and Atlas Integration

In NeuroCaptain v2, the 3D head surface model serves as the primary structural foundation, providing anatomical guidance for both optode montage design and head-cap generation. Extending upon the importing methodologies described in NeuroCaptain 1.0, which were limited to head surface meshes, the workflow now supports 5-layer tetrahedral head models comprising scalp, skull, cerebrospinal fluid (CSF), gray matter and white matter. The scalp and gray matter surfaces are automatically extracted using PyIso2Mesh,^34, 35^ a recent Python port of our MATLAB based 3D surface and volumetric mesh generator. To reduce complexity, only the scalp surface and cortical layers are visualized in the Blender 3D Viewport, while the full volumetric mesh is retained in memory for use in forward light simulations. The scalp surface is used to directly calculate subject-specific 10-20 landmark positions via Brain2Mesh’s `brain1020` function^24, 35^ (now also part of PyIso2Mesh), producing a 10-20 landmark mesh whose vertices correspond directly to the computed landmark coordinates. This landmark mesh serves as the spatial reference for downstream probe registration, enabling optode positions to be encoded as barycentric coordinates relative to landmark vertices and transferred across head surfaces.

In addition to supporting pre-computed surface and tetrahedral mesh models, we further integrate NeuroCaptain v2 with the neuroimaging scans hosted on NeuroJSON.io, a US National Institutes of Health (NIH) funded data portal for sharing standardized neuroimaging datasets. Through this integration, any NeuroJSON.io database containing head surfaces and volumes can be queried and dynamically imported into Blender to be used for montage/head-cap generation. Particularly, the BrainMeshLibrary dataset includes 33 Neurodevelopmental MRI atlases^28, 36^ with ages spanning from 3 months to 85 years. We are currently developing a graphics processing unit (GPU) accelerated full-head segmentation tool, siamize,^37^ based on a deep-learning-based full-head segmentation framework – Segment It All Model (SIAM).^38^ Once siamize is incorporated with NeuroCaptain, one can import from tens of thousands of T1 and T2 weighted MRI scans from NeuroJSON.io and perform segmentation, mesh generation and optode montage/head-cap design in a fully streamlined workflow.

### 2.3 Interactive Optode Montage Design Workflow

Compared to other existing montage design platforms where custom mass-spring models are used, the NeuroCaptain v2 optode montage relaxation model directly leverages the physical simulation engines native to the Blender environment. Optodes (including both sources and detectors) can be added to the head surface near the targeted regions either through automated probe import or interactive placement using intuitive NeuroCaptain panels. Subsequently, each optode is bound to the scalp geometry via the “Shrinkwrap” built-in modifier, which projects the optode position to the nearest point on the head surface and maintains surface normal orientation. When a user interactively drags an optode to reposition it, the Shrinkwrap modifier also ensures the optodes are always constrained to the head surface. Similar to AtlasViewer, a spring-based connection system is used to define the spatial relationships among optodes within the probe. The edges connecting optodes are automatically tessellated based on a user-specified distance threshold. We want to particularly highlight that NeuroCaptain offers capabilities to interactively design modular fNIRS probes – i.e., probes made of repeating modules. Sliding a module spring network over the head surface simultaneously moves all optodes within the module. Optodes designated as anchors are fixed during the relaxation computation, while all non-anchor optodes are free to displace under spring forces until equilibrium is reached.

A 2D schematic of a probe montage designed or registered in NeuroCaptain is generated using an azimuthal equidistant projection of the registered optode, positions, displaying source-detector channel pairs and 10-20 landmark references within the NeuroCaptain panel. Channels are determined by a defined distance threshold. This is rendered within Blender’s image editor, allowing 2D visualization without leaving the application.

### 2.4 Probe Mechanical Constraints and Relaxation Models

NeuroCaptain v2 supports two mechanical relaxation models to accommodate the user’s distance/anchor constraints of the optodes. The first mode is triggered when a user imports a Homer2^39^ and AtlasViewer^25^ compatible “.SD” file. A Python-based spring-relaxation algorithm similar to that implemented in AtlasViewer is invoked to automatically “warp” the 2D probe definition stored in the SD file to the 3D head surface. Anchor labels are matched to the corresponding vertices of the pre-generated 10-20 landmark mesh in the Blender scene, providing the 3D reference positions necessary for registration. Non-anchor optodes are mapped from the 2D probe definition space to 3D head surface coordinates using a per-patch least-squares affine transform fitted from the matched anchor positions, with anchors split into two hemisphere patches to correctly handle bilateral probe layouts, followed by projection onto the head mesh surface. At each relaxation iteration, spring forces are computed for each connection and globally normalized so that the maximum displacement component across all optodes equals one unit. Optode positions are then updated accordingly, and free optodes are re-projected onto the scalp surface. A small time step is used to gradually move non-anchor optode positions until reaching equilibrium. Anchor positions remain fixed throughout relaxation. The SD file defines two categories of springs: stiff springs with explicit rest lengths and flexible springs whose rest lengths are initialized by the inter-optode distance. This allows relaxation to preferentially enforce the geometric constraints encoded in the probe design. This registration procedure is necessary because the SD file encodes optode positions in a 2D probe space that is only weakly linked to the 3D head geometry via the anchor optodes.

The second relaxation model utilizes Blender’s built-in cloth physics solver,^40^ which treats the connection network as a deformable mesh and relaxes it toward the computed rest shape, when importing probe configurations stored in JSON files, unique to NeuroCaptain. Blender’s cloth solver models advanced spring-mass dynamics and is well suited for deforming a probe montage over a curved surface, because optode connections define a network of discrete structural springs. The solver supports four spring types: tension, compression, shear, and angular bending. In our implementation, only tension and compression are enabled, with shear and bending stiffness set to zero, as probe geometry requires only axial stretch and compression resistance along its connections without resistance to in-plane shearing or out-of-plane bending. Each connection in the probe is classified as either “stiff” or “flexible” and carries a pull parameter between zero and one that encodes how strongly that edge (the connection between two optodes) resists deformation.

Because Blender’s cloth simulation does not directly accept per-edge rest lengths, a target rest shape is precomputed before the simulation runs using a simple iterative spring solver that displaces vertices along only the stiff edges toward their per-edge rest lengths defined in the probe configuration. Flexible connections are excluded from this step so they do not impose length constraints on the rest shape. The resulting rest shape defines an optode configuration with each spring at its natural length, representing what the springs “want” in free space, without considering the head surface or anchor constraints.

The cloth simulation then solves the constrained problem on the curved scalp surface. Spring stiffness is modulated per-vertex using vertex groups. Each optode vertex is assigned a weight equal to the mean pull-value of its connected edges divided by 0.9 and clamped to the range [0.01, 1]. Vertices connected to any edge with a defined fixed distance are overridden to a weight of 1.0, ensuring full stiffness for those constraints. Anchor optodes are held in place through the cloth solver’s pinning capabilities: a separate vertex group is created and assigned a weight of one, constraining them to their anatomical coordinate placement. Non-anchor optodes are assigned a weight of 0, allowing them to deform according to the spring forces of the cloth simulation until the forces are balanced.

This relaxation pipeline is evaluated to determine the handling of mechanical constraints based on anatomical variability during initial placement. A probe montage (9 sources, 8 detectors, see Fig. 4a) with 20 stiff springs (high spring constant), 7 flexible springs (low spring constant), fixed-distance links (highest spring constant), and was designed on Colin27 atlas and then registered to two geometrically contrasting NDD atlases (6-0 infant and 35-39 year adult). The deviation of the final relaxed spring lengths from the original rest lengths encoded in the probe configuration file is examined.

### 2.5 Forward Modeling and Cortical Sensitivity Visualization

To further inform and evaluate optode design, quantitative light transport forward modeling and cortical sensitivity visualization are integrated directly into NeuroCaptain v2. Two complementary light transport solvers are supported: the Python binding of our widely used mesh-based Monte Carlo solver (PMMC)^41^ and the diffusion equation (DE) based Redbird^42^ (specifically, its Python port, RedbirdPy). The PMMC solver solves the more general radiative transfer equation (RTE), which is particularly suitable for modeling brain tissues where CSF has low scattering. Redbird, on the other hand, provides an alternative deterministic approach, solving the DE using the finite element method (FEM). Both Python based solvers are directly integrated with the Blender environment and require an imported five-layer head mesh (see Section 2), from which tissue-specific optical properties (absorption coefficient *μ*_*a*_, reduced scattering coefficient *μ*^′^, anisotropy *g* and refractive index *n*) are assigned to each anatomical layer. Simulation parameters are configured via NeuroCaptain panels prior to execution; for PMMC, these include the number of photons and GPU device selection; both solvers accept an optional modulation frequency for frequency-domain probes.

Cortical sensitivity profiles, i.e., the Jacobian with respect to the absorption coefficient, are computed using the adjoint method.^43^ For PMMC, forward simulations are executed independently for each source, followed by adjoint simulations in which each detector is treated as a virtual source. For Redbird, all source and adjoint-source forward simulations are passed to a single batched forward solve, and the sensitivity Jacobian is then computed for all source-detector pairs. In both cases, per-node sensitivity is obtained as the product of the forward and adjoint fluence fields,^43^ summed across all valid source-detector pairs determined by inter-optode distance. The resulting total sensitivity map is visualized directly in Blender by mapping log-scaled sensitivity values to a blue-to-red colormap applied as vertex colors on the cortical surface mesh.

### 2.6 Integration of Optode Montage with NeuroCaptain 3D-Printable Cap Generation

Once an optode montage has been designed or imported within NeuroCaptain v2, the resulting optode locations can be directly integrated into a 3-D printable head cap as built-in grommets that facilitate secure and accurate attachment of the optode or optical fibers. This is enabled by NeuroCaptain’s custom landmark workflow, in which any set of vertices, including one derived from optode positions, may be designated as active landmark placements. Each vertex serves as a cutout placement site during cap generation. A geometry node pipeline projects landmark vertices onto the head surface, instances a cutout shape at each projected location aligned to the local surface normal, and performs a boolean difference to produce cutouts. The geometry and dimensions of each cutout may also be customized, with built-in support for circular, square, and triangular profiles as well as user-supplied geometries, allowing the cap design to accommodate different optode hardware form factors. Apart from this substitution of input positions, the cutout placement and boolean cap generation procedure follows the same workflow described in McCann *et al.*^24^ The standard 10-20 landmark-based workflow remains available, and the choice between the two approaches may be made according to the fabrication requirements of a given study.

### 2.7 Anatomical Coordinates and Cross-Atlas Registration

Instead of using a 2D probe definition that is detached from the 3D head geometry, in NeuroCaptain v2, we use anatomical coordinates to both store and transfer the produced optode montage to a new head model. During export, barycentric coordinates are computed for each optode position relative to the triangulated 10-20 landmark mesh, storing three 10-20 landmark indices, their corresponding anatomical labels, and barycentric weights. These coordinates, along with spring connections and their associated rest lengths, per-connection elasticity parameters, and anchor definitions, are written to a JSON file. Upon import, optode positions are reconstructed from the stored barycentric coordinates relative to the target atlas landmark mesh, providing an anatomically guided initial placement. This approach removes the variability introduced by re-running a spring-relaxation model each time the probe is loaded to a new head model, yielding deterministic results. Following initial placement, optodes are automatically snapped onto the head mesh surface and the spring relaxation is performed to achieve steady-state.

### 2.8 Evaluation of Registration Reproducibility

The variability of optode positions when registering a designed montage over the same head model, and between those from different subjects, is an important performance metric for evaluating a montage design and registration pipeline. For this purpose, we first characterize the reproducibility of NeuroCaptain’s registration methods by quantifying the stability of the anatomical coordinates of the resulting optodes across subjects. We then provide a methodological comparison between two approaches to inter-subject registration: the conventional approach of projecting a two-dimensional optode layout onto a three-dimensional cortical surface, and the approach proposed in this work, i.e., registering optodes directly in 3D space using anatomical coordinates.

For both evaluations, an example probe SD file (“PROBE.sd”) from AtlasViewer is imported into Blender and registered to the Colin27 head atlas.^44^ The registered probe is then saved as an optode JSON file containing the anatomical coordinates and optode connections as well as the mechanical constraints (anchors, spring properties, etc.) for every optode. The exported JSON is then imported and registered onto seven Neurodevelopmental dataset (NDD) young adult head atlases ranging from 18 to 34 years. After cloth-simulation based probe relaxation, the optode positions in the 3D head mesh space are exported, along with the anatomical coordinate weights for every optode. To quantify registration consistency, each subject’s final optode position is projected back onto its registration triangle, yielding three anatomical coordinate weights per subject. The standard deviation of each weight is computed across subjects, and the mean of the three per-optode standard deviations is reported per optode. This metric captures how consistently the optode maintains its relative position within the local landmark geometry, independent of head size.

To further evaluate cross-head-surface probe repeatability, the information from the registered probe is used to compare porting methods. Using the original SD optode file and the NDD atlases, this probe is registered to the seven head surfaces using AtlasViewer. A summary of the inter-subject variability is obtained using the software’s built-in “probe variability” feature, which projects the registered optodes into a common space and takes the mean position and standard deviation for each non-short-separation optode. This process is replicated in NeuroCaptain, where registered optode positions from each atlas are projected into a per-subject Neuromag-style head coordinate frame^31^ constructed from five primary fiducials. The frame origin is defined at the midpoint of the left- (Lpa) and right-preauricular-area (Rpa); the *z*-axis points toward Cz, the *x*-axis points toward the nasion (Nz) and is orthogonalized against *z* via Gram-Schmidt, and the *y*-axis completes a right-handed coordinate system, consistent with the Neuromag MRI coordinate convention. Each axis is then normalized by its corresponding anatomical scale factor, with Nz-Iz applied to the anterior-posterior axis, Lpa-Rpa to the lateral axis, and origin-Cz to the inferior-superior axis, yielding dimensionless coordinates that account for inter-atlas differences in head size and shape. This normalization ensures that positional differences across atlases reflect true anatomical variability in probe placement rather than differences in overall head geometry. Mean position and standard deviation are then computed per optode across atlases in this normalized space, and standard deviations are converted back to millimeters using the mean scale factors across all atlases for reporting. For comparison, only non-short-separation optodes are included to maintain consistency with the AtlasViewer methodology.

The probe variability metrics evaluated here are incorporated into the NeuroCaptain pipeline to enable a quick evaluation method. Each evaluation JSON is exported and saved to a user-defined directory. Once this file has been saved for each head surface, group analysis is automated by reading the evaluation JSON files and generating the per-optode averages and standard deviations as a comma-separated-value (CSV) file.

## 3 Results

### 3.1 Software Interfaces and Optode Visualization

NeuroCaptain renders optode configurations directly on the anatomically derived head surface mesh in Blender’s interactive 3D scene, as illustrated in Fig. 4a. Nine sources and 8 detectors are manually created and added to the scene using the `Add Source` and `Add Detector` functions. The optodes are disc-shaped geometries initially placed at the vertex (Cz) landmark but then seamlessly dragged around the head surface via a “Shrinkwrap” modifier. Sources are rendered as red discs and detectors as black discs for visual distinction. Two optodes designated as anchors are each automatically decorated with a yellow torus indicator, providing unambiguous visual identification of fixed registration points directly in the 3D scene. Optode connections are computed by first projecting the 3D optode positions onto a 2D plane computed by using principal component analysis (PCA), then applying Delaunay triangulation to define the connections between optodes. The resulting edges are added to Blender as a new `Optode Connections` mesh object whose vertices correspond to optode positions and whose edges define the source-detector channel pairs visible in Fig. 4a. The complete probe geometry, including anchor assignments, spring connections, and surface-snapped positions, is designed and verified entirely within the 3D viewport without requiring a separate registration step. A 2D topographic schematic of a probe registered (15 source, 31 detector AtlasViewer example probe) the probe layout is generated using NeuroCaptain and displayed in the Blender space. Sources, detectors and channel pairs, and 10-20 landmarks references are shown in Fig. 4b.

### 3.2 Evaluation of Constraint-Based Spring Relaxation

To evaluate the constraint-based spring relaxation, a NeuroCaptain-designed optode montage (Fig. 4a) is registered and evaluated. A probe originally designed on the Colin27 atlas is exported as a JSON configuration and reimported onto the NDD 30-34 month atlas to demonstrate cross-atlas registration via anatomical coordinate transfer, as illustrated in Fig. 2. All 17 optodes (8 detectors and 9 sources) are successfully placed using the label-based barycentric fallback strategy, as vertex index matching is not available across atlases with differing mesh topologies. The computed positions for the majority of optodes are located in the left-frontocentral region, covered under 10-20 landmarks FCz, FC1, Cz, and C1, consistent with the probe’s intended coverage. Following placement, cloth-simulation based spring relaxation is executed over 100 time steps with a global spring stiffness of 15.0 and gravity disabled, with 2 anchors pinned and 15 non-anchor optodes free to move. The 2 anchor optodes report zero displacement, confirming that pinning is enforced correctly. Non-anchor optodes displace a mean distance of 0.378 mm (range: 0.092–0.820 mm), reflecting convergence of the spring network toward the encoded rest-length geometry on the new atlas surface. Inspection of representative channel pairs shows close agreement between the post-relaxation distances and the JSON-encoded rest lengths: the largest observed deviation is 0.611 mm (Source#5-to-Source#9: current 44.869 mm vs. rest 44.258 mm), corresponding to a relative error of 1.38%, while shorter-range pairs show deviations below 0.5 mm. This validates the expected spring mechanics during the relaxation step of importing a probe montage.

**Fig 2:**
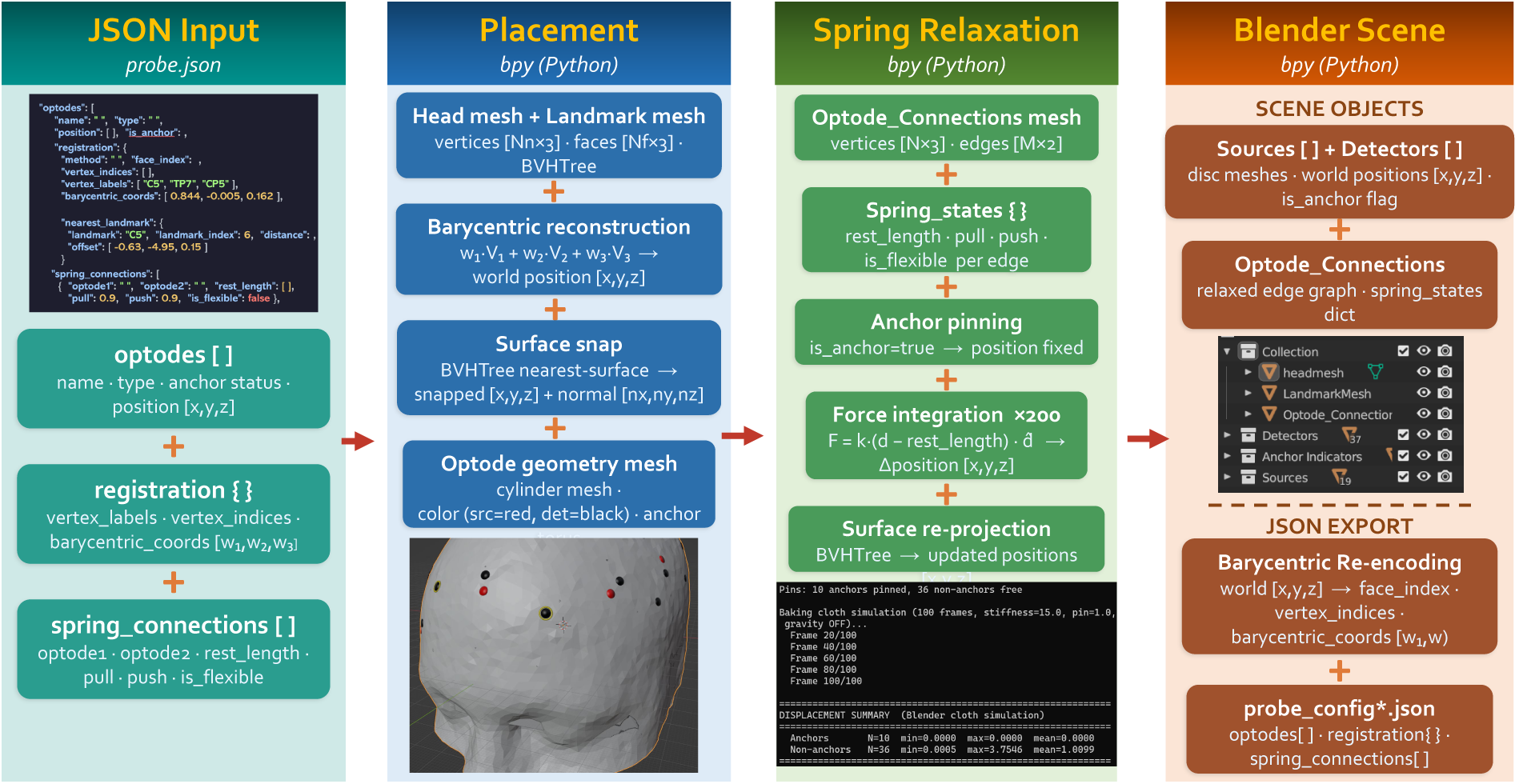
Data exchange and optode porting pipeline in NeuroCaptain v2. The JSON probe format encodes optode identity, barycentric surface coordinates, spring connection parameters, and nearest 10-20 landmark references. This information sets up initial optode placement, and a spring simulation is performed for subtle changes. The probe data can be accessed in Blender for reference and further probe exporting.

**Fig 3:**
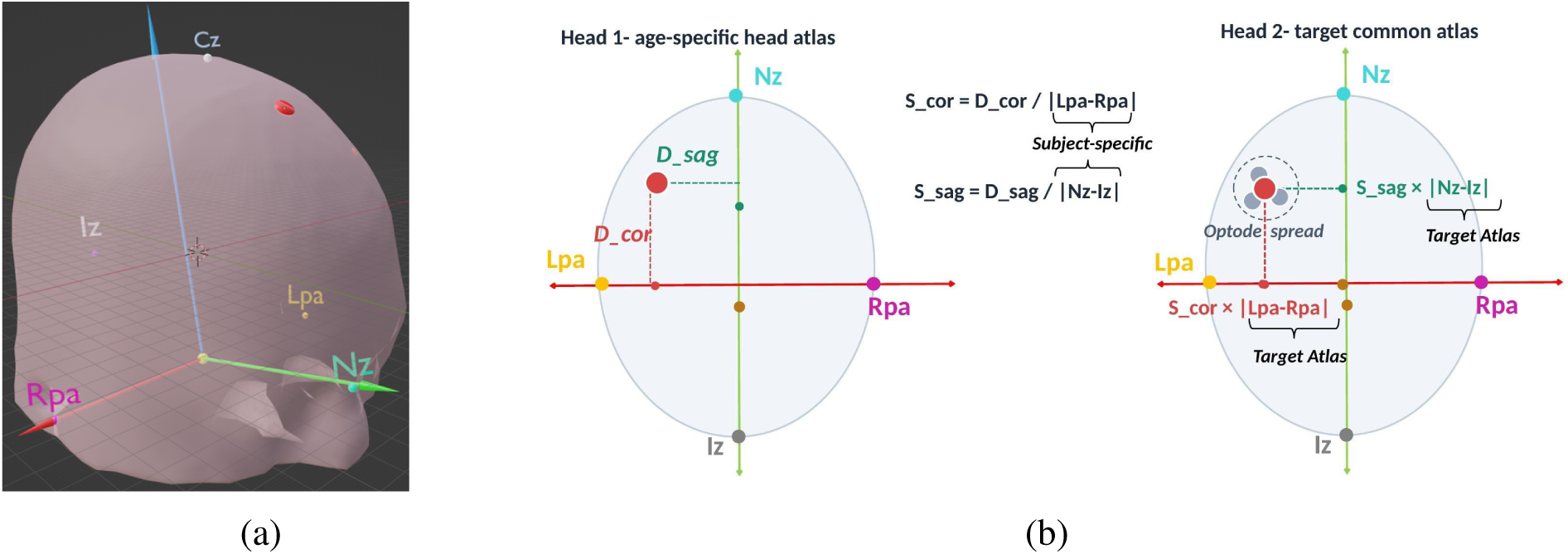
Cross-atlas registration repeatability experiment is guided by the Neuromag MRI coordinate system (a), in which the origin is placed at the midpoint of the pre-auricular fiducials, the *x*-axis points toward the right pre-auricular point (RPA), the *y*-axis points toward the nasion (Nz), and the *z*-axis is defined by their cross product toward the vertex (Cz). Optode positions encoded on a source atlas (Head 1) are normalized by the coronal and sagittal fiducial spans (*S*_cor_ = *D*_cor_/|LPA − RPA|, *S*_sag_ = *D*_sag_/|Nz − Iz|) and re-projected onto a target atlas (Head 2) by rescaling with the corresponding target fiducial distances (b).

**Fig 4:**
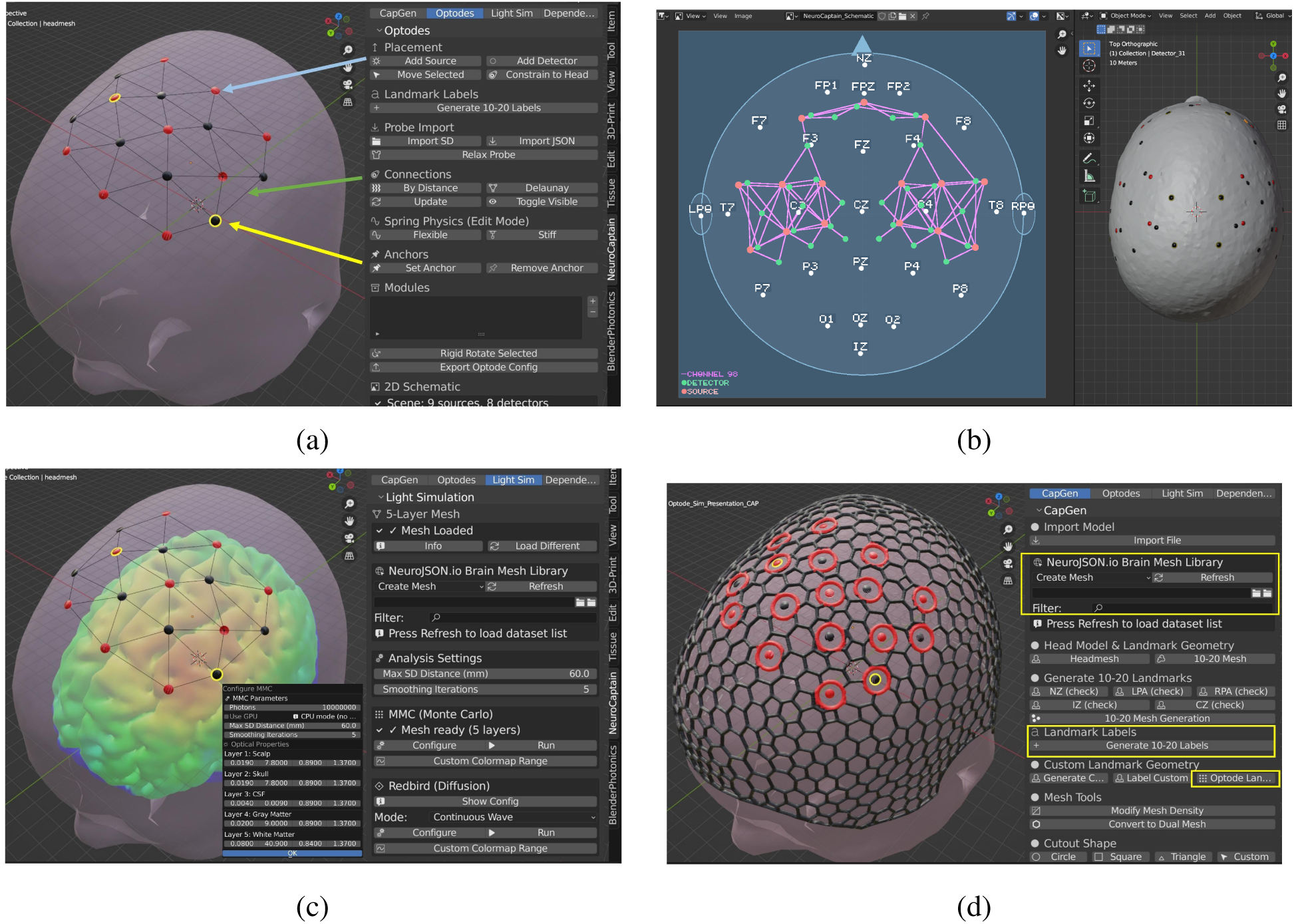
Annotated screenshots of the NeuroCaptain v2 Blender panel are shown. The `Optodes` tab shown in (a) provides controls for manual source and detector placement, spring-based relaxation, anchor assignment, and SD/JSON probe import. This tab also includes a button to generate a 2D probe layout schematic. In (b), NeuroCaptain renders the 2D probe schematic, showing the bilateral sample probe used for registration repeatability tests. The `Light Sim` tab exposes Monte Carlo (MMC) and diffusion-equation (Redbird) solver configuration and launch, together with the NeuroJSON.io brain mesh library for five-layer tetrahedral mesh loading. The result of the PMMC light simulation is visualized in (c). The `CapGen` tab shown in (d) converts the head model 10-20 landmarks, and the optodes into a wireframe 3D-printable cap model.

To evaluate the effectiveness of the cloth-based spring relaxation in accommodating both optode distance constraints and head shape differences, the same probe montage is registered to a 6–0 month infant atlas (NDD 6-0M) and a 35–39 year adult atlas (NDD 35-39). The adult atlas exhibits a 40% larger Lpa–Rpa distance and a 31% longer Nz–Iz distance than the infant atlas, reflecting differences in both head size and shape. After initial placement using the probe anatomical coordinates followed by cloth-based spring relaxation, the stiff springs show a small mean deviation from their original rest lengths of 0.605 mm, which decreases to 0.437 mm for the fixed-distance links (highest stiffness). The flexible springs, which are free to accommodate surface curvature, show a larger mean deviation of 3.724 mm. These results confirm that the cloth-simulation based spring relaxation respects the optode distance constraints while deforming the probe over different head surfaces.

### 3.3 Cortical Sensitivity Visualization

Cortical sensitivity heatmaps computed via forward modeling are rendered directly onto the brain surface mesh, enabling intuitive visual assessment of channel coverage and depth sensitivity. The cortical sensitivity of a probe montage with 9 sources and 8 detectors registered to the scalp surface of a Colin27 5-layer head model, as visualized in Fig. 5a, is evaluated. Fig. 5b demonstrates the configuration of a PMMC simulation, including the tissue-layer optical properties set through the pop-up window triggered by the graphical interface, defining *μ*_*a*_, *μ*^′^, *g*, and *n* values at 630 nm. The photon count is set to 10^7^ and the simulation is run on an NVIDIA GeForce RTX 4090 GPU. The maximum source-detector distance is set to 60 mm, excluding channels that are not sensitive to the cortex.

**Fig 5:**
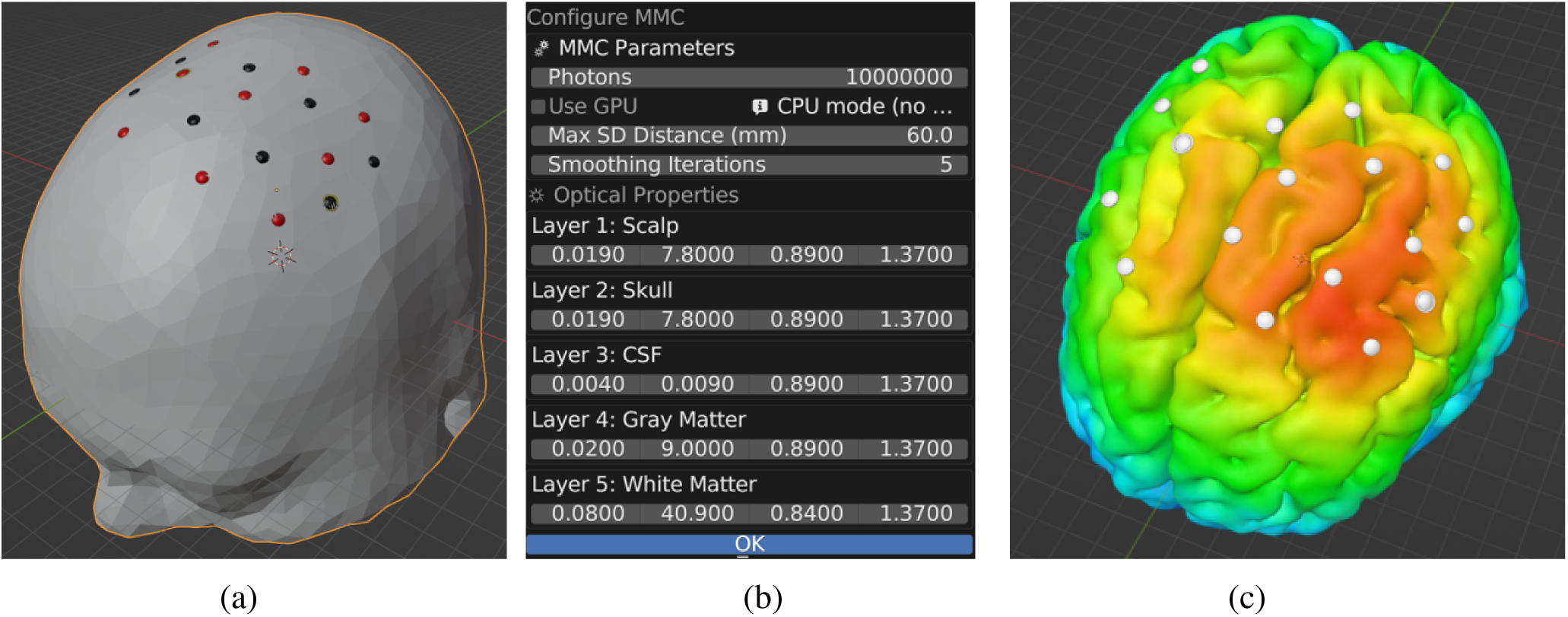
Sample sensitivity map using (a) a registered probe overlaid on the pial surface prior to simulation. MMC solver configuration dialog in (b) is used to customize simulation parameters including: photon count, device selection, and per-layer optical properties for the five-tissue head model. Cortical sensitivity map rendered on the pial surface in (c) following Monte Carlo simulation, with color encoding the logarithmic fluence sensitivity per source-detector channel.

The resulting fluence sensitivity is rendered on the pial surface in the Blender 3D view, as shown in Fig. 5c. The color range is determined by the maximum and minimum sensitivity values computed from the light simulation. The spatial extent of the sensitivity coverage across the cortical surface corresponds to the placement of the probe montage. A Redbird simulation is also performed on the same probe and head model configuration. To improve computational efficiency, mesh cropping is applied using a border of 1 cm to minimize boundary effects. The PMMC simulation launches 10^7^ photon packets using the full Colin27 5-layer head model (402,572 nodes and 2,442,029 tetrahedra). The Redbird simulation uses an optode boundary cropped mesh with a boundary of 1 cm, for a total of 12,781 nodes and 67,900 elements. Run times are measured for the two solvers: 78 s (41 s for forward and 37 s for adjoint Jacobian) for PMMC running on the NVIDIA RTX 4090 GPU and 8 s (forward and adjoint computed combined) for Redbird running on an Intel Core i5-1135G7 processor. Both solvers complete successfully on the same probe and mesh configuration.

### 3.4 Characterization of Registration Repeatability

The repeatability of the NeuroCaptain v2 registration and spring relaxation approach is evaluated using two metrics. First, we characterize the consistency of the anatomical correspondence by computing the per-optode anatomical coordinate standard deviation, obtained by registering the 2D probe defined in the AtlasViewer built-in “PROBE.sd” file across 7 atlas head models. As shown in Fig. 6a, an average standard deviation of the unit-less barycentric coordinates across optodes is *μ* = 0.0159; the lowest variations are reported on the optodes defined as anchors (Sources: 1, 2, 11, 12 and Detectors: 5, 6, 7, 8, 14, 15). This result aligns with the constraints of the registration and spring relaxation, where the anchors have a high pinning force, immobilizing them during this process. We further compute the displacement distances for all optodes between the initial placement after import (*P*_initial_, determined by the saved barycentric coordinates of the probe using

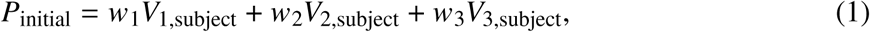

where *w*_*i*_ and *V*_*i*_ denote the barycentric coordinates (weights, 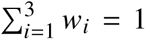) and the adjacent 10-20 landmark positions, respectively, and the final positions after cloth-simulation based spring relaxation. An average displacement of 0.33 mm is reported, indicating high positional stability within the triangle encoded by the anatomical coordinates. It should be noted that this metric captures intra-triangle consistency only and does not reflect absolute optode placement variation across atlases, which is separately quantified below.

**Fig 6:**
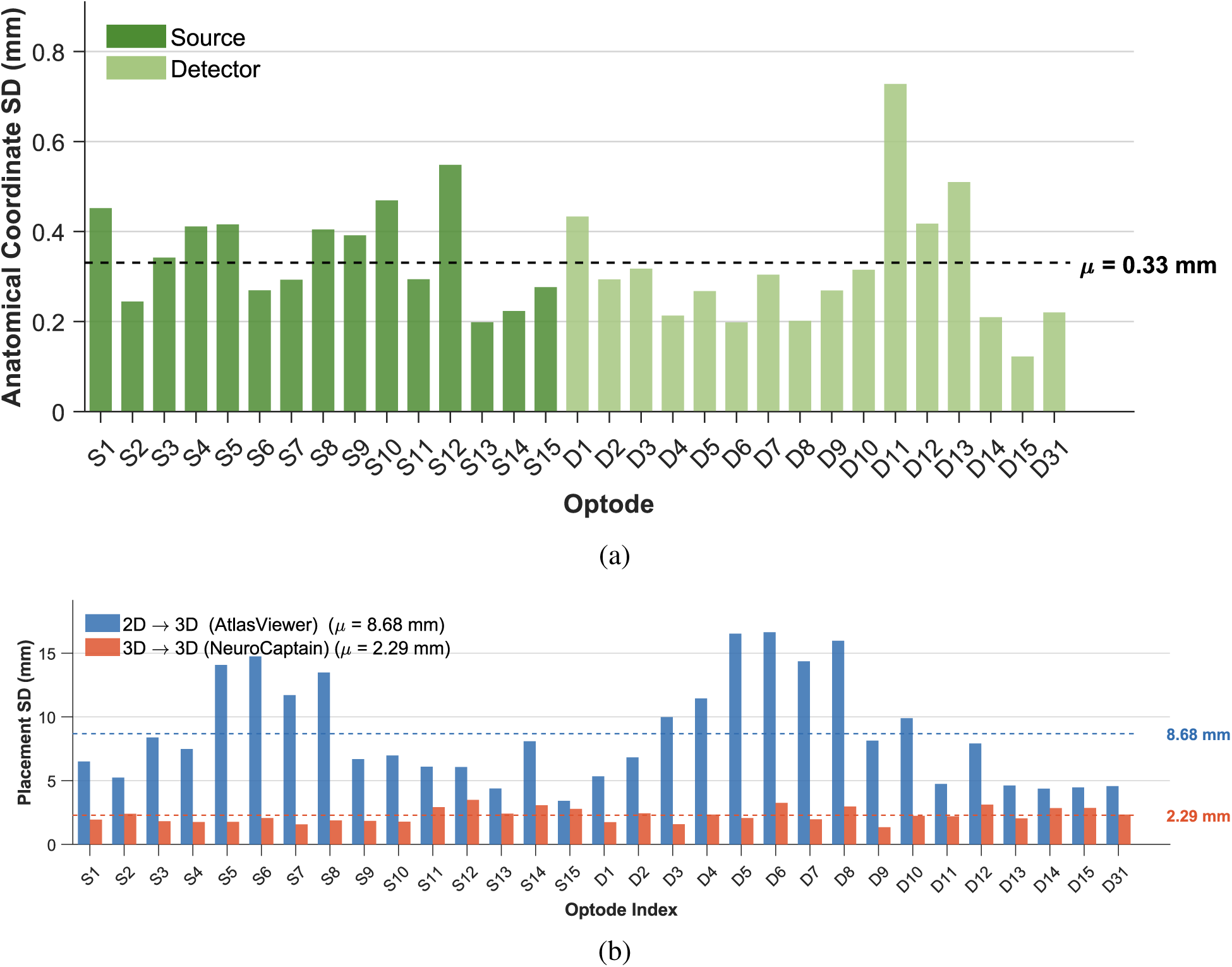
Assessment of registration repeatability of a benchmark probe across 7 head models. The average standard deviation of each barycentric weight is visualized in (a) for each optode in the probe, with dark green bars corresponding to sources (S1-S15) and light green bars corresponding to detectors (D1-D15, D31). A horizontal dotted line is used to visualize the average of 0.0159. To further evaluate the performance of using anatomical coordinates, we compare with a 2D probe to 3D registration in (b). After transforming the optodes from 7 different atlases to a common space, the standard deviation for each method is reported, with blue bars corresponding to 2D →3D and orange corresponds to anatomical coordinate registration. There is an ∼74% reduction in the average variability.

Registration of probe configurations across head atlases of varying shapes reveals the cross-subject reproducibility achievable with the barycentric anchor registration approach.

Global optode registration variability is determined by projecting the optodes into a common anatomical space and examining the standard deviation of each optode. This metric compares two registration methods: the anatomical coordinate approach proposed in this paper (3D → 3D) and a conventional 2D → 3D registration. The latter method produces a mean optode standard deviation of 8.68 mm across all 7 atlases, whereas the anatomical coordinate registration approach reduces the average optode standard deviation to 2.29 mm – an approximately 74% reduction in variability, as depicted in Fig. 6b.

## 4 Discussion

Probe montage design and analysis is a crucial component of neuroimaging studies, directly impacting cortical coverage and subsequent analyses. NeuroCaptain v2 introduces several new tools to aid in optode montage design, probe registration and sensitivity analysis. The technologies we have developed and validated include streamlined access to diverse head mesh libraries via the NeuroJSON.io data portal, real-time 3D interactive probe design, utilization of Blender’s cloth-simulation based spring relaxation, storage and transfer of designed optodes using anatomical coordinates, and integration of optode montages into 3D-printable cap designs.

As illustrated in the screen captures in Fig. 4, real-time 3D optode montage design provides an intuitive platform that places optodes in a manner directly comparable to the experimental implementation. Designing in 3D removes the need for an iterative design approach, as the montage can be visualized in the same configuration in which the probe is ultimately used. The optodes are placed into the Blender space with simple buttons on the graphical user interface (GUI). Optode connections can be defined by either Delaunay triangulation or by distance. These connections can be customized as long as the resulting connection mesh is named `Optode Connections`, the name that the subsequent functions reference when they require this information. This allows flexibility in the variety of montages that can be designed in NeuroCaptain, including modular designs. Anchors can also be easily defined and are visualized with a yellow torus so that they remain visually distinctive during the design process.

Implementing Blender’s cloth solvers allows us to tailor our registration approach to the proposed anatomical coordinate registration. While this initial registration shape preserves the anatomical correspondence of the original optode montage design, inter-optode distances can change with varying head shape and size, so the structural constraints are violated. Blender’s cloth solver provides per-edge spring classification, which controls which edges participate in the rest-shape computation that encodes target lengths from the probe design, separate from the initial on-scalp placement. Per-vertex stiffness modulation controls how strongly each optode resists deformation during cloth simulation, as the springs find equilibrium freely with a final projection to the surface, decoupling the surface projection used for the SD import. This relaxation method uses a sophisticated solver to balance the structural constraints of the probe montage against its anatomical placement.

Visualizing an optode montage’s sensitivity to the cortical surface is essential for ensuring proper ROI coverage, and, when computed across multiple subjects, consistent coverage across individuals can also be evaluated. Implementing both PMMC and Redbird allows users to select a solver based on their accuracy requirements, project stage (prototyping vs. final), and computational and hardware resources. These solvers are easily configurable using the pop-up window and are automatically visualized in the same space as the optode design, enabling an easy transition between design and evaluation.

A simple yet effective approach that NeuroCaptain v2 introduces is the migration from the traditional 2D flat/Cartesian representation of a probe to a 10-20 anatomical-coordinate based representation. The porting of optodes uses anatomical coordinates to encode the anatomical information associated with each optode position. By storing optode locations as barycentric coordinates within the tessellated 10-20 landmark mesh, the probe definition is tied to the anatomical geometry of the head rather than to a particular coordinate on the mesh topology. This allows probe montages to be transferred across head models without a 2D-to-3D projection, which would otherwise introduce large distortions and additional sources of variation, and provides a deterministic initial placement. The combination of anatomically grounded initialization and subsequent cloth-based relaxation simultaneously addresses two distinct challenges of probe porting: preserving anatomical correspondence and satisfying mechanical constraints.

The effectiveness of the NeuroCaptain montage design and cross-subject registration pipeline is reflected in the registration reproducibility results shown in Figs. 6a-6b. The mean barycentric standard deviation across 7 atlases is 0.0159 (unit-less), corresponding to an average positional stability of 0.33 mm within the triangle encompassing each optode. This demonstrates that the anatomical coordinate representation is stable across atlases of varying head geometries, even after probe relaxation. At the global level, projecting registered optode positions into a common anatomical frame revealed a mean optode standard deviation of 2.29 mm using the anatomical coordinate registration approach, compared to 8.68 mm using 2D-to-3D registration, demonstrating a 74% reduction in variability. The underlying reason for this improvement is that the anatomical coordinate registration initializes each optode at an anatomically consistent location across atlases, rather than re-deriving 3D positions from a 2D layout or using 3D world coordinates, which may map inconsistently across head surfaces.

Once a probe montage has been designed, its optode positions can be directly integrated into a 3D-printable head cap as built-in grommets. This integration leverages the custom workflow described in McCann *et al.*,^24^ and the cap generation procedure follows the same pipeline as the original NeuroCaptain, with the optode positions substituted in place of the standard 10-20 landmarks. Compared to NinjaCap^23^, which also produces 3D-printable caps that can embed optode grommets, NeuroCaptain’s integrated optode montage design workflow greatly simplifies this process. The ability to carry a validated optode layout through to a fabrication-ready cap design without leaving the NeuroCaptain environment reduces the risk of introducing errors during format conversion or manual reentry of optode positions.

We would like to note a few limitations of this current study. The selection of cranial landmarks (Nz, Iz, Lpa, Rpa, and Cz) from the head surface remains a manual step and the accuracy of the resulting 10-20 points depends on this selection. The current evaluation uses a set of seven NDD young adult atlases, so broader validation across atlases representing a wider range of ages, head sizes, and ethnic origins would further characterize the generalization of the approach. An important consideration in probe design is the balance between mechanical constraints and anatomical placement: over-constraining a probe, for example, by assigning fixed rest lengths to too many connections, may prevent the spring relaxation from adapting adequately to a new head geometry, while under-constraining may allow optodes to drift from their intended anatomical targets. While the probe variability feature provides quantitative feedback to help users identify and address such issues, interpreting these metrics and adjusting spring parameters accordingly may not always be intuitive. While the forward modeling integration supports interactive sensitivity evaluation, the computational cost of PMMC, even with GPU acceleration, may still limit truly real-time iterative probe adjustment; accelerating light simulation to more closely approach real-time feedback remains an important direction for future work. Future extensions may also benefit from rendering the cortical parcellations according to standard atlas labels, enabling sensitivity to be reported directly in terms of coverage of specific anatomical ROIs rather than as an aggregate map.

## 5 Conclusion

In summary, we present NeuroCaptain v2, a comprehensive and fully open-source workflow for fNIRS optode montage design, cross-subject registration, and cortical sensitivity visualization, implemented as an easy-to-use add-on to the widely adopted Blender 3D modeling platform. By allowing researchers to directly add, edit, and optimize source and detector positions over a 3D head surface, NeuroCaptain v2 circumvents the indirect 2D-to-3D iterations required by conventional montage design tools and unifies probe definition, performance evaluation, and head-cap fabrication within a single anatomical environment. The designed montage is stored as barycentric coordinates within a tessellated 10-20 landmark mesh, tying the probe definition to the underlying head anatomy rather than to a particular mesh topology, and enabling deterministic, anatomically guided probe transfer across diverse head models. Combined with Blender’s cloth-based spring relaxation, this representation simultaneously preserves anatomical correspondence and satisfies the mechanical constraints of the probe, reducing the cross-atlas registration variability by approximately 74% relative to a conventional 2D-to-3D approach. By integrating our model-based forward simulators, the mesh-based Monte Carlo (MMC) solver and the diffusion-equation solver Redbird, NeuroCaptain v2 further offers direct, quantitative guidance for sensitivity-informed optode placement. The workflow capitalizes upon streamlined access to a wide variety of head models hosted on the NeuroJSON.io data portal, as well as the rich 3D modeling capabilities of Blender. We anticipate that the wide accessibility of this open-source tool will help the community standardize fNIRS probe montage design, improve cross-lab reproducibility, and streamline the path from anatomically informed probe design to fabrication-ready head caps.

## Data and Code Availability

The NeuroCaptain v2 Blender add-on is open-source and can be freely accessed at https://github.com/COTILab/NeuroCaptain.

## Acknowledgments

This research is supported by the National Institutes of Health (NIH) National Institute of Neurological Disorders and Stroke (NINDS) grant U24-NS124027, National Cancer Institute grant R01-CA204443, and National Institute of General Medical Sciences (NIGMS) grant R01-GM114365. Agentic coding assistant Claude Code (Anthropic, CA) was used for assisting the development of the NeuroCaptain v2 software; Claude was also used for proofreading the manuscript.

Biographies and photographs of the other authors are not available.

